# Changes in Sound Localization Performance of Single-Sided Deaf Listeners after Visual Feedback Training in Azimuth

**DOI:** 10.1101/2020.04.18.048363

**Authors:** Bahram Zonooz, A. John Van Opstal

## Abstract

Chronic single-sided deaf (CSSD) listeners lack the availability of binaural difference cues to localize sounds in the horizontal plane. Hence, for directional hearing they have to rely on different types of monaural cues: loudness perceived in their hearing ear, which is affected in a systematic way by the acoustic head shadow, on spectral cues provided by the low-pass filtering characteristic of the head, and on high-frequency spectral-shape cues from the pinna of their hearing ear. Presumably, these cues are differentially weighted against prior assumptions on the properties of sound sources in the environment. The rules guiding this weighting process are not well understood. In this preliminary study, we trained three CSSD listeners to localize a fixed intensity, high-pass filtered sound source at ten locations in the horizontal plane with visual feedback. After training, we compared their localization performance to sounds with different intensities, presented in the two-dimensional frontal hemifield to their pre-training results. We show that the training had rapidly readjusted the contributions of monaural cues and internal priors, which resulted to be imposed by the multisensory information provided during the training. We compare the results with the strategies found for the acute monaural hearing condition of normal-hearing listeners, described in an earlier study [1].

## Introduction

The healthy auditory system applies three types of acoustic cues to localize a sound [2]. For sound sources in the horizontal plane (azimuth angle), the system uses interaural differences in arrival times (ITDs, order: up to ±600 *µs*), and intensity (ILDs, up to about ±20 dB). The latter arises from the head-shadow effect (HSE), which causes frequency-dependent sound attenuations: the higher the frequency, the stronger the shadowing by the head, and the left-right difference can thus provide a unique frequency-dependent cue for source azimuth. Below approximately 1.5 kHz, the HSE is too small to be reliably detected by the brain (< 1 dB), but for these lower frequencies the interaural time/phase differences become a reliable azimuth cue. Note that for all locations in the midsagittal plane the ITDs and ILDs are zero, and therefore cannot specify the elevation direction of a sound source. The elevation angle can be extracted from the complex broad-band spectral shape cues that result from direction-dependent reflections and refraction of high frequency (> 4 kHz) sound waves within the pinna cavities, and can be characterized by direction-dependent head-related transfer functions (HRTFs; [2, 3, 4, 5, 6, 7]).

The auditory system needs to map the implicit acoustic localization cues to veridical twodimensional sound-source directions in azimuth and elevation to achieve a coherent and accurate percept of sound location. These cues change in the course of one’s life-span, which suggests that the auditory system should be able to recalibrate and reweight those cues as the head and ears slowly grow and change. In earlier studies from our lab [8, 9], we have shown that the capacity to relearn new cues is not only limited to early childhood. For example, the auditory system can adapt to chronic [10] and acutely-imposed changes of the pinnae [8, 11, 12, 9].

We recently showed that normal-hearing human listeners rapidly learn to remap the acoustic spectral cues for elevation during a short training session to a limited number of locations, both with and without visual feedback [13]. In a follow-up study [1], we subsequently demonstrated that the auditory system can also swiftly reweight its internal priors regarding the binaural difference cues and monaural head-shadow cues, to improve localization performance in azimuth, in response to acutely imposed monaural hearing (ear-canal plug and muff over one ear).

While there is consensus that rapid adaptation is possible in the auditory system [14, 15, 16, 8, 17, 18, 19], so far, studies have reported different results on the reweighting of the different monaural and binaural localization cues and prior information sources to estimate a sound’s direction [20, 21, 22, 23].

In this paper, we set out to study short-term adaptive behavior of chronic single sided deaf (CSSD) listeners, and compare the results with acute conductive unilateral plugged (ACUP) listening for normal-hearing subjects, as described in the previous study [1]. In the latter group, the manipulation perturbed the highly robust binaural differences cues for sound-source azimuth, which led to an immediate and dramatic degradation of sound-localization performance in the horizontal plane, with a large localization bias towards the hearing ear. Interestingly, the deficit in binaural hearing, however, also impaired localization in the vertical plane, i.e., up-down and front-back directions, as binaural spectral integration for the elevation angle is known to be mediated by perceived azimuth [12, 24], which in the case of (acute) binaural impairment is erroneous. Interestingly, after learning with visual feedback, localization performance in azimuth improved, which could be partly attributed to a recalibration of the perturbed binaural differences, but for elevation it deteriorated. Possibly, learning to also use the (monaural) spectral cues to estimate azimuth, may have interfered with the capacity to use the same cues for elevation. Alternatively, the listeners could have adopted a stronger weight for a near-zero elevation prior in response to the feedback learning.

A similar problem might be present in single-sided deaf listeners, albeit manifested differently, as Chronic single-sided deaf listeners cannot rely on any binaural difference cues to localize sounds in the horizontal plane. Furthermore, as Chronic single-sided deaf listeners may have had ample time to (potentially) adapt to their monaural listening condition, they may have learned to employ different localization- and cue-weighting strategies than acutely plugged normal-hearing listeners who lack such long-term monaural experience.

As illustrated in Figure 1, we hypothesize that Chronic single-sided deaf listeners could potentially make use of three monaural acoustic cues for directional hearing in the horizontal plane (see Appendix, for further details): (i) the overall acoustic head shadow, which changes the perceived (broadband) sound level at the hearing ear in a systematic (sinusoidal) way with source azimuth (e.g., [25]; Eq. A1), (ii) low-pass spectral filtering of the head, which causes an azimuth-dependent spectral head shadow, in which higher frequencies are attenuated more than lower frequencies, creating an azimuth-dependent effective bandwidth at the hearing ear, and (iii) monaural spectral-shape cues from the pinna of the hearing ear, in which higher-order spectral-shape features, like notch width, steepness, and depth, could provide unique azimuth-dependent information.

**Figure 1:**
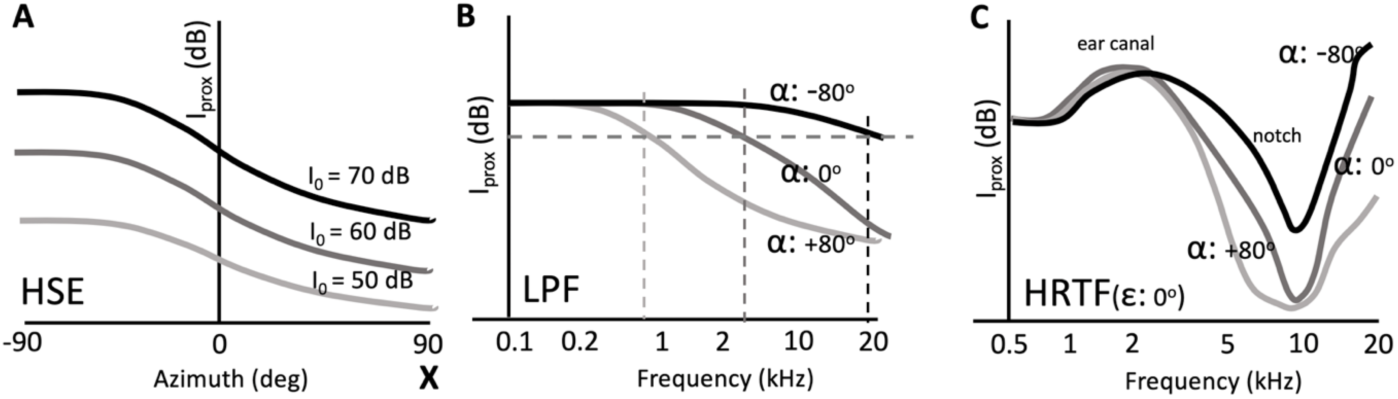
Three potential monaural head-shadow cues that are available to single-sided deaf listeners. Each cue varies with source azimuth in a different way. (A) The overall proximal sound level (integrated across all frequencies) varies with azimuth and absolute sound intensity (here indicated for 50, 60 and 70 dB; Eqn. A1). X indicates the deaf side (right ear). (B) The head’s main spectral effect is approximated by a low-pass filter. The cut-off frequency (vertical dashed lines) for a flat spectral source (at fixed intensity) varies with source azimuth, *α* (Eqn. A5). (C) The spectral fine structure of the HRTF (at elevation zero deg) also varies with source azimuth, as high frequencies are attenuated more than low frequencies. This leads to shape changes (widening and deepening) of high-frequency notches and peaks.

Note, however, that each of these cues is in principle ambiguous, as infinitely many sound-source spectra and azimuth combinations can yield the same acoustic input at the hearing ear. As such, monaural sound localization is inherently ill-posed. Therefore, to deal with this problem, the listener should combine the acoustic sensory cues with additional sources of prior information, e.g. regarding the expected absolute source intensity, potential locations, and potential spectral profiles.

To investigate their short-term adaptation behavior and to compare their results with those obtained from acute conductive unilateral plugged subjects, three Chronic single-sided deaf listeners were trained, with visual feedback about the true sound location, to localize a sound of fixed spectral content and intensity at only a limited number of locations in the horizontal plane. Listeners generated a fast head-orienting saccade to the perceived sound location, as well as a fast corrective head movement to the visual feedback stimulus. After training, they were tested for their localization behavior in the entire frontal hemifield. We discuss the preliminary results from these Chronic single-sided deaf listeners, by comparing their adaptive behaviour with the results from acutely monauralized listeners.

## Materials and Methods

### Participants

Three single-sided deaf listeners (M1-M3, ages M1: 25, M2: 53, and M3: 59; 1 female) participated in the free-field sound-localization experiments. All were naive regarding the purpose of the study. M1 and M3 were deaf in their left ear, M2 in the right ear. The non-affected ear of all three listeners had normal hearing, within 20 dB HL (see Table 1). Subjects were given a brief practice session to get acquainted with the setup and localization paradigm, and to gain stable localization performance to standard broadband Gaussian white noise stimuli.

**Table 1:** Standard audiograms measured across 5 octaves from 0.25°8 kHz for the three listeners. All three have normal hearing in their good ear, and severe sensorineural hearing loss (> 80 dB) in their impaired ear.

| SUBJ | Ear | 0.25 | 0.5 | 1.0 | 2.0 | 4.0 | 8.0 | kHz |
| --- | --- | --- | --- | --- | --- | --- | --- | --- |
| M1 | **Left** | 85 | 100 | 100 | 100 | 100 | 100 | dB |
|  | **Right** | 15 | 10 | 5 | 5 | -5 | 0 | dB |
| M2 | **Left** | 5 | 5 | 10 | 15 | 25 | 20 | dB |
|  | **Right** | 110 | 120 | 120 | 120 | 110 | 110 | dB |
| M3 | **Left** | 110 | 120 | 120 | 120 | 120 | 110 | dB |
|  | **Right** | 20 | 20 | 10 | 25 | 15 | 25 | dB |

### Ethics statement

The local Ethics Committee of the Faculty of Social Sciences of the Radboud University (ECSW, 2016) approved the experimental procedures, as they concerned non-invasive observational experiments with healthy adult human subjects. Prior to their participation in the experiments, all subjects gave their full written consent.

### Experimental setup

During the experiments, subjects sat comfortably in a chair in the centre of a completely dark, sound-attenuated room (length x width x height: 3 × 3 × 3 m). The walls of the 3 × 3 × 3 m room were covered with black foam that prevented echoes for frequencies exceeding 500 Hz. The background noise level in the room was about 30 dB SPL [26].Target locations and head movement responses were transformed to double-polar coordinates [27]. In this system, azimuth, α, is defined as the angle between the sound source or response location, the center of the head, and the midsagittal plane, and elevation, ε, is defined as the angle between the sound source, the center of the head, and the horizontal plane. The origin of the coordinate system corresponds to the straight-ahead speaker location. Head movements were recorded with the magnetic search-coil induction technique [28]. To that end, the participant wore a lightweight (150 g) “helmet” consisting of two perpendicular 4 cm wide straps that could be adjusted to fit around the participant’s head without interfering with the ears. On top of this helmet, a small coil was attached. From the left side of the helmet, a 40 cm long, thin, aluminum rod protruded forward with a dim (0.15 Cd/*m*^2^) red LED attached to its end, which could be positioned in front of the listener’s eyes, and served as an eye-fixed head pointer for the perceived sound locations. Two orthogonal pairs of 2.45 2.45 m coils were attached to the edges of the room to generate the horizontal (60 kHz) and vertical (80 kHz) magnetic fields. The head-coil signals were amplified and demodulated (Remmel Labs, Ashland, MA), after being low-pass filtered at 150 Hz (model 3343; Krohn-Hite, Brockton, MA) before being stored on hard disk at a sampling rate of 500 Hz per channel for off-line analysis.

### Auditory Stimuli

Acoustic stimuli were digitally generated using Tucker-Davis Technologies (TDT) (Alachua, FL) System III hardware, with a TDT DA1 16-bit digital-to-analog converter (50 kHz sampling rate). A TDT PA4 programmable attenuator controlled sound level, after which the stimuli were passed to the TDT HB6 buffer and finally to one of the speakers in the experimental room. All acoustic stimuli were derived from a standard Gaussian white noise stimulus, which had 0.5 ms sine-squared onset and offset ramps. This broadband GWN control stimulus had a flat characteristics between 0.2 and 20 kHz, and a duration of 150 ms. The three types of stimuli were presented during the experiments. Broadband (BB), Low-pass (LP) and High-pass (HP) contained the frequencies from 0.2 to 20 kHz, all frequencies up to 3.0 kHz and the frequencies above 3.0 kHz, respectively (Figure 2). In the adaptation experiment, only the HP stimuli were chosen as by focusing on the HP stimuli we excluded the ITD contribution to azimuth sound localization. Absolute free-field sound levels were measured at the position of the listener’s head with a calibrated sound amplifier and microphone (Bruel and Kjaer, Norcross, GA).

**Figure 2:**
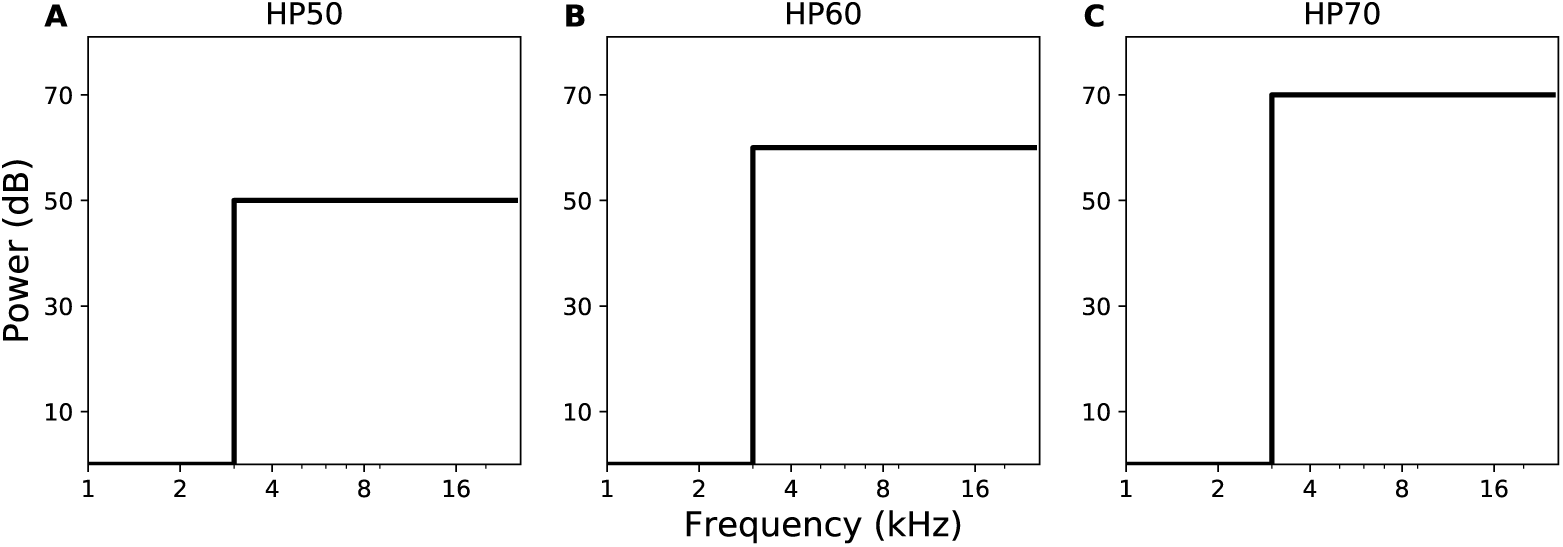
Schematized power spectra of the sound stimuli used in the pre/post-adaptation experiment. Stimuli were derived from a GWN control stimulus by (A) removing all frequencies below 6 kHz (HP) with (A) 50 dB (B) 60 dB (C) 70 dB.

### Experimental paradigms

#### Calibration

Each experimental session started with a calibration paradigm to establish the mapping parameters of the search-coil signals to known target locations. Head-position data for the calibration procedure were obtained by instructing the listener to make an accurate head movement while redirecting the dim LED in front of the eyes from the central fixation LED to each of 58 peripheral LEDs, which was illuminated as soon as the fixation point extinguished. The 58 fixation points and raw head-position signals thus obtained were used to train two three-layer neural networks (one for azimuth, one for elevation) that served to calibrate the head-position data, using the Bayesian regularization implementation of the back-propagation algorithm (MatLab; Neural Networks Toolbox) to avoid overfitting [29]. In each sound-localization experiment, the listener started a trial by fixating the central LED (azimuth and elevation both zero; Figure 3, [1]). After a pseudo-random period between 1.5°2.0 sec, the fixation LED was extinguished, and an auditory stimulus was presented 400 msec later. The listener was asked to redirect the head by pointing the dim LED at the end of the aluminum rod to the perceived location of the sound stimulus as fast and as accurately as possible.

**Figure 3:**
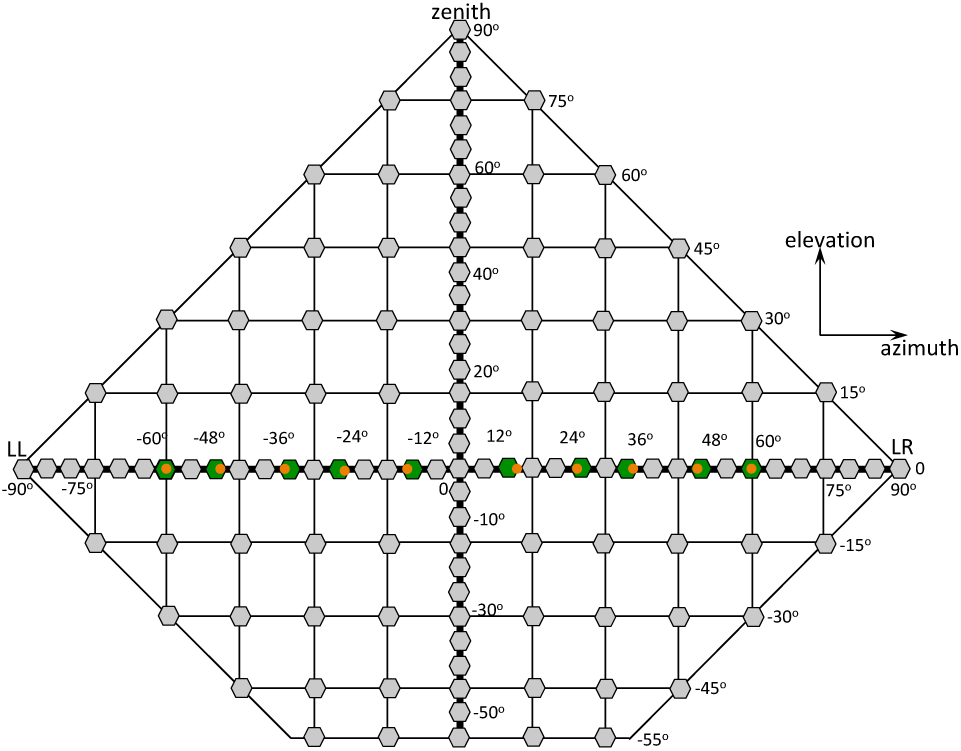
Distribution of sound-source locations, as used in the different experimental paradigms, projected in a flattened Cartesian azimuth-elevation coordinate grid. Note that speakers were attached to a spherical frame, and that in the double-pole azimuth-elevation coordinate system the sum of azimuth and elevation angles can never exceed 90 deg (outer diamond-like boundary of the plot). The training targets were located on the azimuth plane, and are indicated in red. They were presented with visual feedback (green dot) at the end of each trial. The pre- and post-adaptation test targets (red and dark grey) were distributed across the frontal hemifield, and were pseudo-randomly selected for azimuth in [−60,60] deg, and for elevation in [−40,+50] deg. In the control experiment of day 1, the selected speaker locations were confined to [−60,+60] deg for azimuth, and [−40,+40] deg for elevation. LL: lateral left, LR: lateral right. The central speaker at (0,0) deg, and the speaker at the zenith were not used [1].

#### Control session

The sound-localization experiments were carried out over two experimental days. The localization control experiment was performed on the first day. This experiment contained 300 trails with broadband, low-pass and high-pass stimuli, and were presented at randomly selected locations that ranged from [−60,+60] deg in azimuth, and from [−40,+40] deg in elevation (see Figure 3). To prevent successful use of the HSE [25], the stimuli varied in intensity; sound levels of the HP stimuli varied between 45 dB and 70 dB in 5 dB increments; sound levels of the LP and BB stimuli were either 50 dB or 65 dB (HP: 6 different sound levels, 30 locations, in total: 180 trials, and LP, BB each 2 different sound levels, 30 locations, in total 120 trials). The control experiment served to establish the subject’s localization abilities, and to assess the effect of sound level on the monaural listeners’ localization performance, prior to the adaptation experiment. That is, we chose a sound level for which they had a considerable bias toward the normal-hearing ear. The pre-adaptation, training, and post-adaptation experiments were performed on a second recording day.

#### Training

In the training experiment, subjects localized the HP stimuli at 60 dB, presented at 10 fixed locations in the azimuth direction (+60, +48, +36, +24, +12, −12, −24, −36, −48, −60 deg), at elevation zero (Figure 3). After the sound was presented, and the subject had made the localization response, a green LED in the center of the speaker was illuminated for a duration of 1500 ms. The subject had to make a subsequent head-orienting response to the location of the LED; this procedure ensured that the subject had access to signals related to programming a corrective response, immediately after the sound-localization estimate. The training experiment consisted of 400 trials in which every location was presented 50 times in pseudo-random order.

#### Test sessions

The pre- and post-adaptation experiments contained the same 180 trials, consisting of three types of stimuli: HP50, HP60, and HP70 sounds. Stimuli were presented at pseudo-randomly selected locations in the full 2D frontal hemifield, ranging from [−60,60] in azimuth, and from [−40, +50] deg in elevation.

## Data Analysis

We analyzed the calibrated responses from each participant, separately for the different stimulus types, by determining the optimal linear fits for the stimulus–response relationships for the azimuth and elevation components:

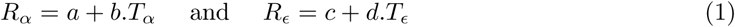

by minimizing the least-squares error, using the Scikit-learn library. *R*_*α*_ and *R*_*ϵ*_ are the azimuth and elevation response components, and *T*_*α*_ and *T*_*ϵ*_ are the actual azimuth and elevation coordinates of the target. Fit parameters, a and c, are the response biases (offsets; in deg), whereas b and d are the response gains (slopes, dimensionless) for the azimuth and elevation response components, respectively. Note that an ideal localizer should yield gains of 1.0, and offsets of 0.0 deg. We also calculated Pearson’s linear correlation coefficient, *r*, the coefficient of determination, *r*^2^, the mean absolute residual error (standard deviation around the fitted line), and the mean absolute localization error of each fit. Linear regression for listener M2 were performed on the inverted azimuth coordinates of the stimulus–response relations, in order to align the deaf side to the left (positive bias) for all listeners.

### Multiple linear regression

To test to what extent the acute monaural listener makes use of the ambiguous head shadow effect (HSE) and the true source location (presumably through distorted remaining binaural cues, and spectral cues, see above) to localize sound sources, we analyzed our data with a multiple linear regression. We evaluated the relative contributions of sound level and stimulus azimuth to the subject’s azimuth localization response in the following way:

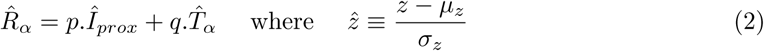

Here, 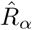, 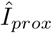, and 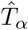 are the dimensionless z-scores for the response, proximal sound level, and target values, respectively, with *µ*_*z*_ the mean, and *σ*_*z*_ the standard deviation of variable z. In this way, the contributions of sound level and sound location can be directly compared, although they are expressed in different units, and may cover very different numerical ranges. The partial correlation coefficients, p and q, quantify the relative contributions of sound level and target azimuth, respectively, to the measured response. An ideal localizer would yield *p* = 0 and *q* = 1, indicating that the localization response is not affected by variations in perceived sound level. On the other hand, if *p* = 1 and *q* = 0 the responses are entirely determined by the head-shadow effect. The proximal sound level, 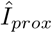, was calculated as the perceived intensity at the free ear, by using the following approximation:

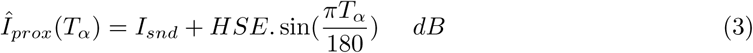

Here, *I*_*snd*_ is the actual free-field sound level (in dBA) at the position of the head, and the sine function approximates the head-shadow effect and ear-canal amplification for a broad-band sound (we took *HSE* = 10 dB; see [27]).

For the elevation responses, we extended the multiple regression analysis in the following way [25]:

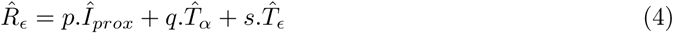

Here, the elevation response was considered to potentially depend on proximal sound level, the true target’s azimuth location, and the true target’s elevation angle. For an ideal localizer, the partial correlations should yield [*p, q, s*] = [0, 0, 1].

## Results

### Azimuth

#### Controls

Listeners were first exposed to control experiments in which elicited goal-directed head saccades to ten different sounds. Figure 4 shows the results for listener M2 in azimuth for different stimuli. Responses were highly inaccurate for all three stimuli, as gains and biases substantially deviated from the optimal values of 1.0 and 0.0 deg, respectively. The positive slopes of the regression lines were, however, significantly different from zero, and varied from one stimulus to the next. This indicates that the stimuli still appeared to contain some valid localization cues. Importantly, the localization bias changed in a systematic way with sound level for all three spectral stimulus types. As this listener is deaf in the left ear, soft sounds were perceived mainly towards the deaf side (negative bias) while the louder sounds shifted toward the hearing side (hearing ear). This result differs from the acute conductive unilateral plugged listeners who displayed a large hearing-ear bias for all sounds in this experiment.

**Figure 4:**
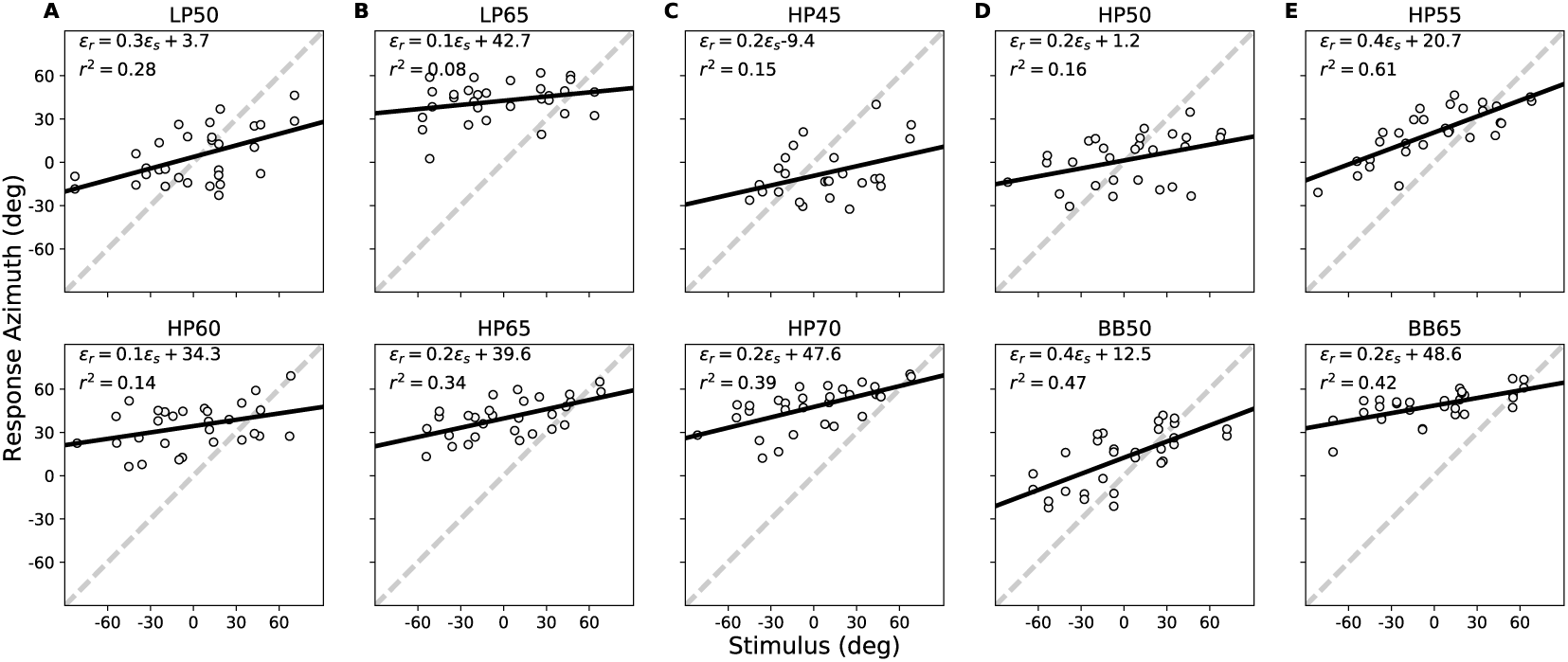
Localization results for listener M2 in azimuth for the ten control stimuli (each point corresponds to a single trial). Responses were highly inaccurate, as the gains and biases deviated substantially from their optimal values of 1.0 and 0.0, respectively. Note the systematic change in the localization bias with sound level, which is apparent for the LP, HP and BB sounds. The listener perceived soft sounds mainly towards the deaf side, or to the central space (negative or near-zero bias), and loud sounds on the hearing side (large positive bias).

#### Pre-training

On the second day of the experiments, listeners performed the localization task for three high-pass filtered stimuli at different levels (HP50, HP60 and HP70). Figure 5 shows the pre-training regression results in azimuth for listener M2. The data indicate poor localization performance as both gain and bias were far from the optimal values. However, it is quite clear that the bias increased for higher sound levels: the HP50 stimui yielded smaller response biases (−9.3 deg) than the higher intensity stimuli, HP60 (18.2 deg) and HP70 (47.2 deg). Also, the response gains were significantly larger than zero.

**Figure 5:**
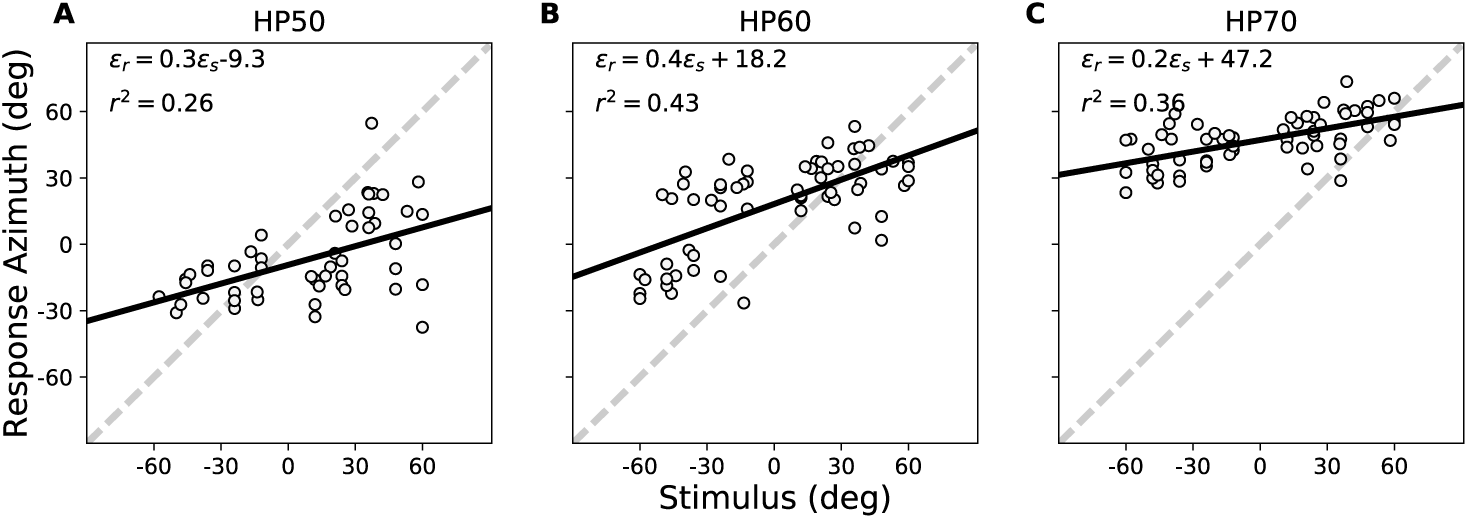
Pre-adaptation: localization results for subject M2 in azimuth for the three test stimuli, prior to the training session. Responses were highly inaccurate, as the gains and biases deviated substantially from their optimal values of 1.0 and 0.0, respectively. Yet, the slopes of the regression lines differed significantly from zero, indicating that the stimuli contained some azimuth-dependent localization cues.

#### Training

In the training experiments, listeners were exposed to 401 trials in which they had to respond with a saccadic head movement to a HP60 stimulus, randomly selected out of ten locations in the azimuth plane. Immediately after the first localization response, an LED was presented at the center of the speaker to provide visual feedback about the actual source location. The subjects were instructed to make a corrective head movement towards the LED. This experiment was done to check whether visual feedback would lead to improvements in the localization performance. Figure 6 shows the regression results of listener M2 during the course of the training session for three windows of 50 trials: at the beginning of the training session (trials 1-51), after the first phase of the training (trials 101-151), and near the end of the training (trials 351-401). Looking at the gain and bias obtained for the three windows, it is clear that both values improved as training advanced: the gain increased from *b* = 0.5 to 0.7, while at the same time the bias reduced from *a* = 12.1 to 1.2 deg. At the same time, the response precession increased from *r*^2^ = 0.6 to 0.8.

**Figure 6:**
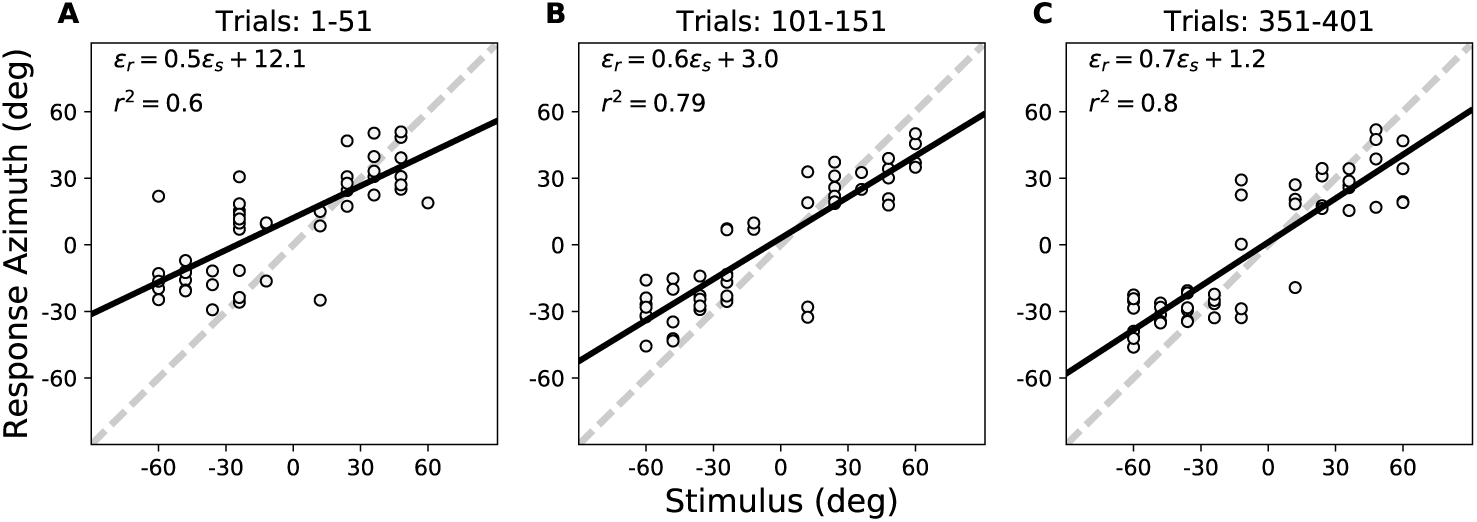
localization data of M2 for the ten training targets (HP60 stimuli) presented in randomized order with visual feedback in the azimuth plane (elevation zero) at the start (trials 1-51), after 100 training trials (nrs. 101-151), and towards the end of the session (trials 351-401). Note the systematic increase of the response gain, and the reduction in response variability (increased *r*^2^) and bias during the session (cf. Figure 5B).

#### Post-training

Immediately after the training session, subjects had to perform the same experiment as the pre-training session, without the visual feedback, to investigate whether their localization performance differed from the pre-training results. Figure 7 shows the regression results for M2. Comparingn the stimulus-response plots for this listener with the pre-training performance (Figure 5), reveals that the training had clearly affected localization performance, not only for the limited set of ten trained locations, but also for non-trained locations, and for the different stimulus levels. The gain for stimulus HP60 had increased from *b* = 0.4 to 0.6, the bias decreased by nearly 25 deg towards the deaf side (from pre: *a* = +18.2 deg, down to post: *a* = −6.5 deg), while at the same time localization precision improved as well (*r*^2^ increased from 0.43 to 0.61). The behavior for the lower (HP50) and higher (HP70) intensity stimuli, however, appeared to be different. For these sounds, the response gains remained low, at 0.2 − 0.3, but the biases were more strongly expressed: the soft sounds were heard more into the deaf side (bias decreased from *a* = −9 to −40 deg), whereas the louder sound remained well at the hearing side (bias changed from *a* = +47 to +31 deg).

**Figure 7:**
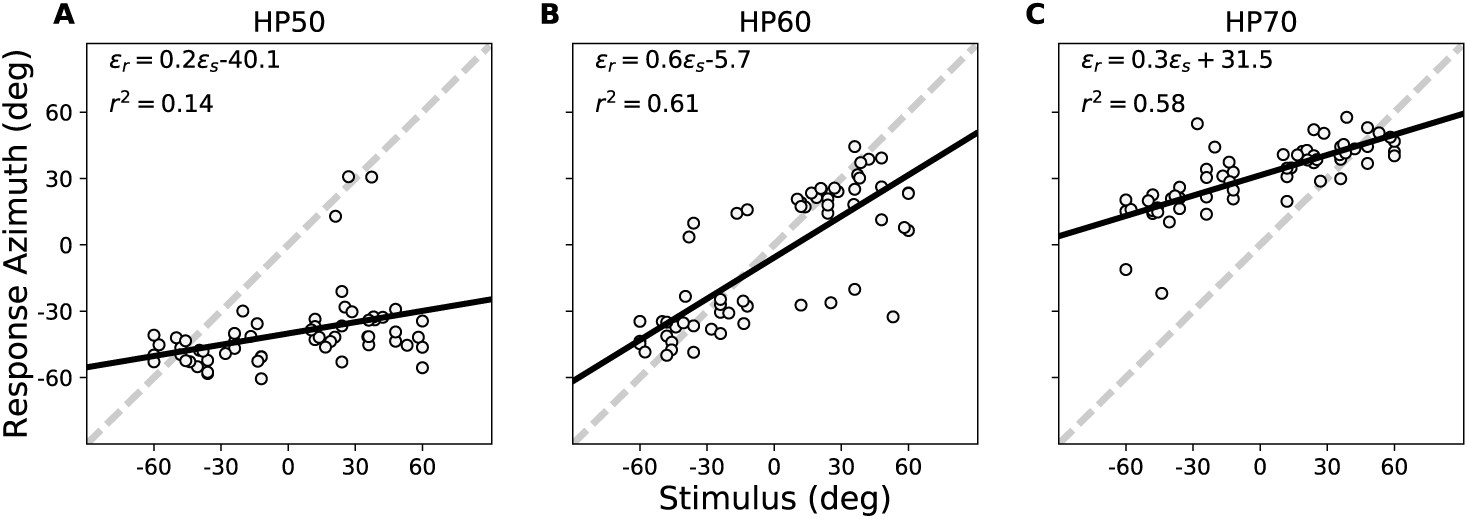
Stimulus-response plots for the azimuth components for M2 immediately after adaptation. Comparison of these data with Figure 5 shows that training changed localization performance for the non-trained azimuth-elevation locations and stimulus levels considerably: the bias for the 60 dB sound is now close to zero, with an increased gain, whereas the softer sounds were localized far into the deaf hemifield, and the loudest sound remained far into the hearing side, both with the same low gain.

To illustrate the training effect for the three listeners, we summarized the overall statistical results for pre- and post training sessions in Figure 8 for the HP50, HP60, and HP70 stimuli before and after training. The most consistent improvements were obtained for the trained HP60 stimuli: increased gains and precisions, lowered biases and MAE’s. The results for the softer sounds showed no change in gain, *r*^2^ and MAE (data points near the diagonal), and a significant increase in the negative bias, as sounds were consistently localized more into the deaf side. A slight localization improvement was observed for the higher-intensity sounds, as the MAE was significantly reduced, because of the reduced bias. These response patterns, albeit preliminary, seem to differ markedly from the results obtained with the acute unilateral plugged hearing condition described in [1].

**Figure 8:**
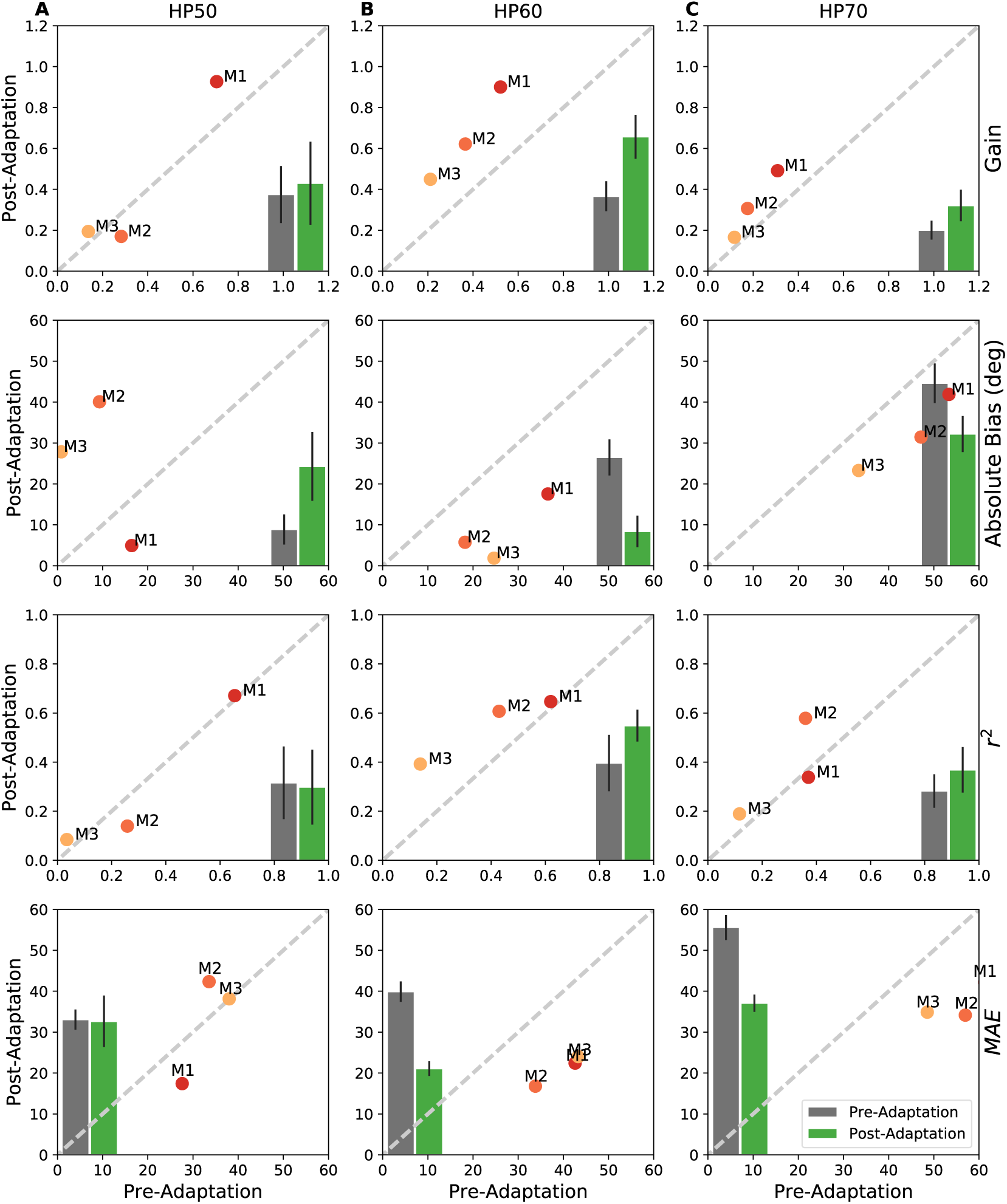
Summarized statistics of regression analysis for the three participants for pre-adaptation and post-adaptation. Columns: the three different test stimuli; top row: response gain, center row: response bias; third row: coefficient of determination, and bottom row: mean absolute error (in deg). Averages across listeners are shown as insets: grey = pre-adaptation mean with std, green = post-adaptation mean with std. For the HP60 sounds, the post-adaptation results are more accurate (higher gains, and smaller bias), and more precise (less variability, higher *r*^2^). The HP70 stimuli (right) yielded smaller overall errors, because of the reduced positive bias. Gains and precisions, however, did not change. The HP50 stimuli only yielded larger negative biases (into the deaf side), except for M1, who had a high response gain for this sound.

To describe better how subjects had changed their localization performance we also performed a multiple regression analysis (Eqs. 2 and 3) on the pre- and post-adaptation data to quantify the contributions of the HSE and the true azimuth location to the responses. Figure 7B shows that the HSE (indicated by proximal sound level) had a large (negative) contribution (around −0.70) to the responses, but did not change significantly with visual feedback training. In contrast, the partial correlation coefficients for the true azimuth locations had increased significantly from 0.78 ± 0.09 to 0.90 ± 0.11 for the three listeners (Figure 9C).

**Figure 9:**
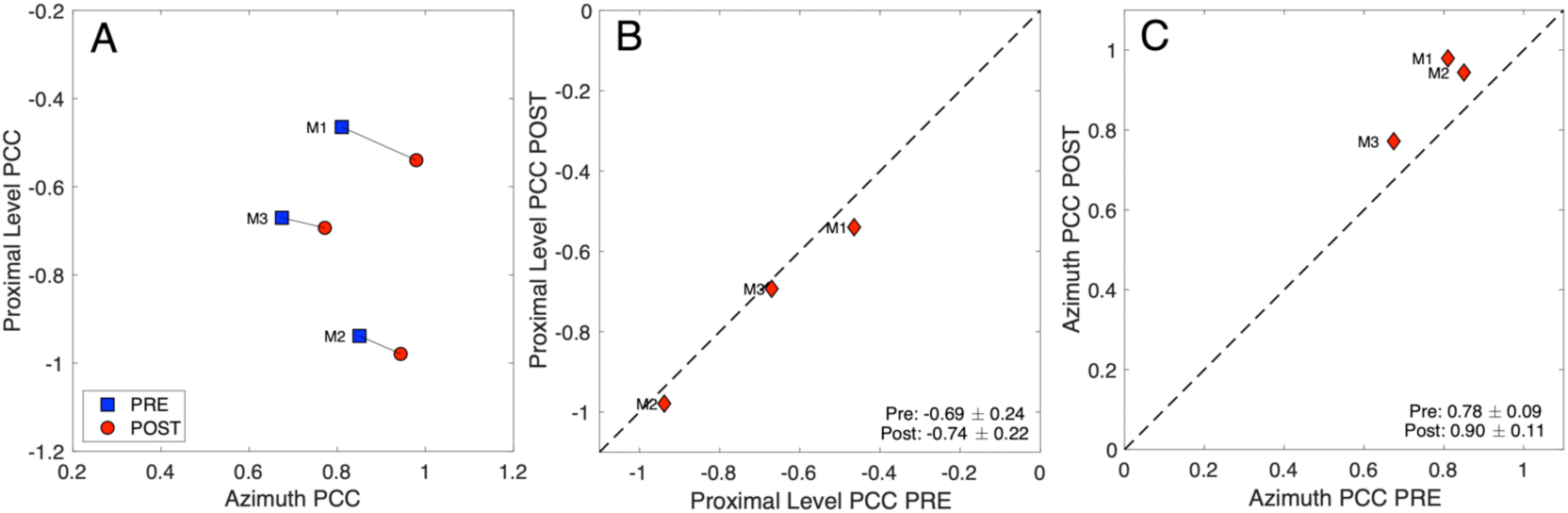
Multiple linear regression results for the three participants. The contribution of proximal sound level to their responses is high, but did not change with visual-feedback training (B). The azimuth component, however, did increase for all three listeners, by approximately the same relative amount (C).

## Elevation

To assess the potential training effect on the elevation performance of the SSD listeners, we performed multiple linear regression according to Eqn. 4. Figure 10 shows the results for this analysis. The partial correlation coefficients of HSE to the elevation responses did not show significant change with visual feedback training. The data however indicate that the partial correlation coefficients of true azimuth locations increased for all three subjects, while the elevation contribution remained unchanged.

**Figure 10:**
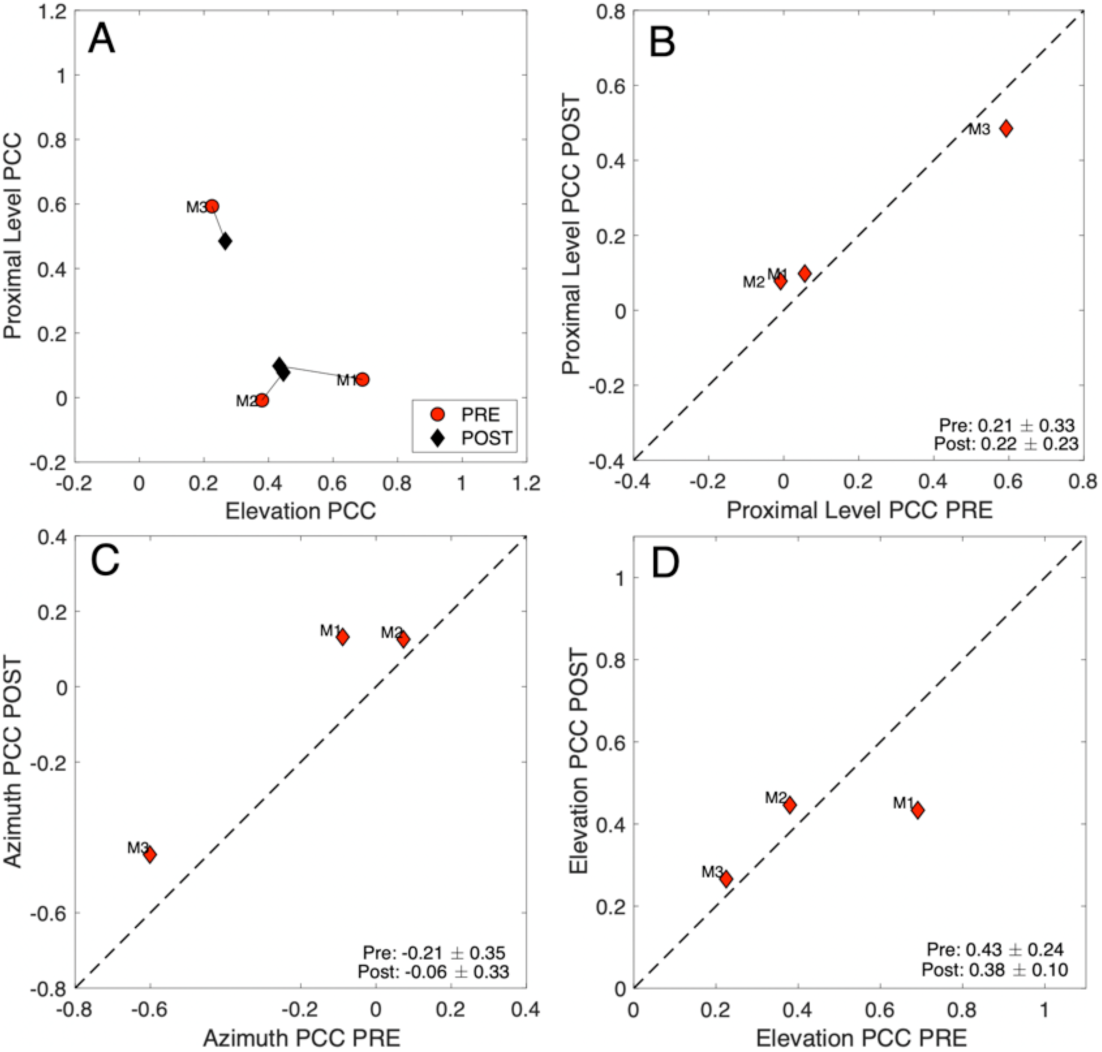
Multiple linear regression (Eqn. 4) on the elevation responses for the three SSD participants. The pre-training contribution of proximal sound level to the elevation responses did not change significantly with visual-feedback training (B). The azimuth component increased for all three listeners (C), whereas the elevation contribution remained unaffected, except for M1, whose elevation PCC had decreased after training (D).

## Discussion

In addition to the acoustic information, there are also types of non-acoustic information, such as prior knowledge, attention, or memory of specific non-acoustic details related to sound source, that brain uses to localizes sounds. For example, it is highly unlikely that the sound of a car originated from above or below. A normal binaural listener can localize sounds accurately based on the measured acoustic information. Such a normal hearing listener will, therefore, put a smaller weight on the non-acoustic information than the acoustic information. These non-acoustic information are even more important for Chronic single-sided deaf. However, The mechanisms underlying adaptive processes to Chronic single-sided deaf are still unclear. Our experiment thus set out to establish the process in which the learning the system had to cope with ongoing changes. We also compared the short-term adaptation behavior of Chronic single-sided deaf listeners to that of acute conductive unilateral plugged listener.

To study the mechanisms underlying the integration of the different acoustic cues, and the effect of adaptation on the chronic monaural listeners, they were presented with sound sources. We studied the adaptation in the Chronic single-sided deaf azimuth localization system. Our experiment thus set out to establish the process in which the auditory system had to cope with ongoing changes. We carried out the experiments where the subjects were exposed to training session for a fixed-intensity high pass sound source, presented at limited number of locations in the horizontal space. They had to generate head-orienting responses to the sound sources distributed in a set of restricted locations in azimuth plane. By providing visual feedback during the training session we investigated the ability of the listeners to cope with sound sources of various levels and spectra. Each trial was followed by the exact same sound source with an LED pointing to the sound location. The subjects, therefore, could correct head movement errors trial after a trial.

We observed from Figure 6 that during the training Chronic single-sided deaf improved her localization performance towards the end of the training for HP60. Our results also showed that the adaptation generalized to other target locations, and to the intensity 70, indicating that adaptation was not a simple cognitive trick, however, in acute conductive unilateral plugged listener adaptation generalized to all sound intensities. Unlike acute conductive unilateral plugged listeners, the adaptation in the horizontal direction in Chronic single-sided deaf did not affect the elevation responses This can indicate that in Chronic single-sided deaf spectral pinna cues might underlie the improved performance in horizontal plane. This is perhaps due to the fact that Chronic single-sided deafs have developed a mechanism during their lives that increased use of pinna cues from hearing ear to azimuth direction does not interfere with the estimation process for elevation. All three Chronic single-sided deaf subjects slightly increased HSE effect during training. In the pre-adaptation sound localization behavior for acute conductive unilateral plugged, azimuth direction is mainly determined by ITDs, monaural intensity, filter cues from HSE, and potential weakened ILDs. For Chronic single-sided deaf, however, our data suggest that sound level plays a strong role with a week contribution of low pass spectral filtering, as there is no contribution from surviving ILDs at all. The post-training localization data indicate that Chronic single-sided deaf subjects put A larger weight on the good-ear HRTF which led to improved azimuth performance, while the elevation estimate is modulated by the azimuth precept (Figure 11). However, for acute conductive unilateral plugged subjects after training, the elevation localization resulted to be highly determined by the prior.

**Figure 11:**
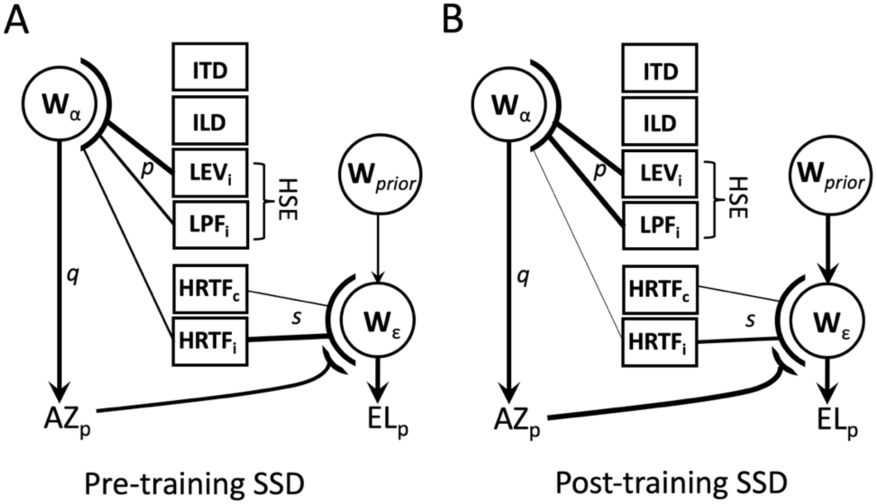
Potential available localization cues for SSD listeners. (A) Because of the absence of any contralateral input from the deaf ear, only three monaural cues remain to estimate source azimuth (sound level, the good-ear HRTF and the low-pass head filter). The pre-adaptation data suggest a strong contribution of Iprox and a weaker contribution from azimuth (from ipsi-spectral or LP filter cues). The elevation percept could be based on azimuth and HRTF cues, in combination with prior assumptions. (B) After training, the azimuth-related weights increased for azimuth estimation, possibly from increased weights of HRTF and low-pass head filter, whereas the elevation percept is based on a stronger azimuth percept. In listener M1, also the prior weight had increased.

## Acknowledgements

Supported by the European Union Program FP7-PEOPLE-2013-ITN ’HealthPAC’, Nr. 604063IDP (BZ), Horizon 2020 ERC Advanced Grant ’Orient’, Nr. 693400 (AJvO). We would like to thank the subjects who volunteered to participate in the experiments.

## Author Contributions

BZ and AJVO designed research; BZ performed the study; BZ and AJVO analyzed data; BZ and AJVO wrote the paper.

## Appendix Three potential monaural localization cues for azimuth

### Monaural Cue 1

A simple acoustic model for the overall head-shadow effect, illustrated in Figure 1A (after [25]), holds that for a flat broad-band sound of intensity *I*_0_, the proximal sound intensity at the (right) hearing ear can be approximated by:

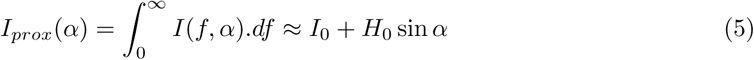

where *H*_0_ is the maximum head-shadow effect (approximately 10 dB for broad-band noise); the plus sign indicates that the right ear is the hearing ear. For the left ear, this becomes a minus sign.

### Monaural Cue 2

The head effectively acts as an azimuth-dependent low-pass filter (Figure 1B). We thus assume that the cut-off frequency is azimuth dependent, given as *f*_*c*_(α), so that a flat BB sound with intensity *I*_0_ will be filtered as:

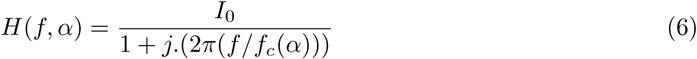

Its amplitude characteristic is calculated as:

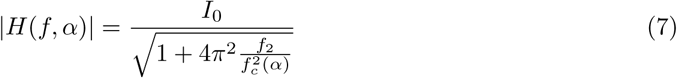

and the total power, up to the cut-off frequency (i.e., the LP head shadow) is then:

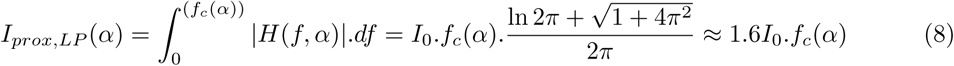

which varies linearly with the cut-off frequency (= effective bandwidth) and absolute sound level. Suppose that for the cut-off frequency we can take the following heuristic approximation: it is maximum (say, 19 kHz) when the sound is presented at the far free ear (at azimuth −90 deg) and approaches its minimum, say 1.0 kHz, when the azimuth is at the opposite side, at +90 deg (the deaf side). A simple function that fulfills this requirement is

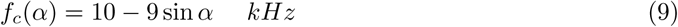

Combined with Eq. 8, this gives

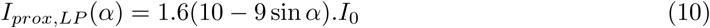

which is very similar to Eqn. 5, although it is based on a different principle (spectral analysis).

### Monaural Cue 3

Analysis of the spectral cues, illustrated in Fig. 1C, would require some sort of spectral pattern recognition algorithm on stored monaural HRTFs. This could be done in a similar way as has been proposed for the elevation angle (e.g. []Hofman and Van Opstal, 1998), for which it has been suggested that the auditory system may perform a spectral correlation analysis. The incoming sensory spectrum (computed from the convolution between the source and the actual source direction) is compared with stored representations of all HRTFs. Then, either maximum-likelihood estimation (which detects the elevation angle for which the spectral correlation is maximal; Hofman and Van Opstal, 1998), or a Bayesian estimate that includes an elevation prior to yield a posterior distribution (e.g., []Zonooz et al., 2019) would uniquely point at the veridical (or optimal) elevation angle, provided that the source spectrum itself is uncorrelated with any of the stored HRTFs.

New here is that a similar algorithm could in principle be applied to extract both the azimuth and the elevation angle, when assuming (as schematically indicated in Figure 1C) that the changes in azimuth would lead to consistent (and learnable) changes in e.g. the shape, steepness, width, and/or depth of the different high-frequency peaks and notches, caused by the low-pass filtering characteristics of the head (Figure 1B).

